# A novel periplasmic layer formed by an outer membrane lipo-protein governs the cell-envelope integrity and stiffness of *Leptospira interrogans*

**DOI:** 10.64898/2026.02.11.705417

**Authors:** Keigo Abe, Nobuo Koizumi, Hiroko Takazaki, Mika Hirose, Kyosuke Takabe, Takayuki Kato, Shuichi Nakamura

## Abstract

The structural integrity and mechanical properties of the cell envelope are crucial for bacterial physiology and dynamics; however, how specific molecular components govern these properties remains poorly understood. Here, we report the novel structural and mechanical functions of LipL32, the most abundant outer-membrane (OM) lipoprotein in the pathogenic spirochete *Leptospira interrogans*. LipL32 accounts for approximately 75% of the total OM proteins, yet its functional role has remained enigmatic. Cryo-electron microscopy revealed that the highly dense LipL32 forms a hitherto unappreciated thin layer in the periplasmic space, positioned adjacent to the peptidoglycan (PG) layer. The structural analysis also demonstrates that the disruption of *lipL32* by transposon insertion results in irregular deformation of OM and spatial variability in the width of the periplasmic space. Quantitative measurements by optical tweezers confirmed that the *lipL32::*Tn mutant exhibited a significant reduction in whole-cell stiffness. Consistent with this mechanical impairment, the *lipL32::*Tn mutant displayed a diminished ability to penetrate a heterogeneous agar mesh model that mimics the host’s extracellular matrix, suggesting an important role for cell rigidity in initial infection via damaged skin. These findings define LipL32 as a critical mechanical stabilizer that maintains the structural integrity of the cell envelope. This work provides a new paradigm for understanding how specialized lipoproteins contribute to bacterial cell mechanics, offering insights into pathogenic invasiveness.

## Introduction

The cell envelope of Gram-negative bacteria is more than a passive barrier; it is a complex, multi-functional organelle that governs the cellular mechanical properties and physiology (1), and is associated with various dynamics such as motility (2). Regarding the outer membrane (OM), its role as a permeability barrier is well-established, but recent studies revealed a variety of functions heretofore unknown. For instance, though the peptidoglycan (PG) has been recognized as a major determinant for cell shape, OM is also involved in the improvement of bacterial cell shape through modification by the enhanced synthesis of lipopolysaccharides (LPS) (3). Also, the connection between the OM and PG via a small α-helical OM lipoprotein, called Braun’s lipoprotein (Lpp), confers resistance against osmotic shock on *Escherichia coli* (4).

Among Gram-negative bacteria, spirochetes, including pathogens such as *Borrelia burgdorferi* (Lyme disease), *Treponema pallidum* (syphilis), *Brachyspira hyodysenteriae* (swine dysentery), and *Leptospira interrogans* (leptospirosis), are structurally distinguished by the presence of periplasmic flagella (PFs) situated between the OM and PG layer (5). While the core machinery for OM biogenesis and protein secretion in spirochetes is largely conserved with other Gram-negative bacteria, their OM components are enriched with diverse proteins involved in host interactions and immune evasion (6). For instance, OspA is a major immunogenic lipoprotein in *B. burgdorferi* (7). DbpA and DbpB of *B. burgdorferi* function as adhesins with binding affinity for the extracellular matrix (ECM) component decorin (8), while *T. pallidum* Tp0751 exhibits affinity for laminin (9).

This study focuses on the OM proteins of pathogenic *Leptospira*, the causative agent of leptospirosis. Leptospirosis is a worldwide zoonosis, exhibiting an extremely wide range of host spectra (10). *Leptospira* spp. possesses two short PFs (one flagellum per cell end) in the short-pitch helical cell body and moves in liquid and on surfaces (Fig. 1) (5, 11). *Leptospira* spp. comprises pathogenic and nonpathogenic (saprophytic) species. The pathogenic strains are colonized in the proximal renal tubules of maintenance hosts, typically rodents in nature, and shed into the environment upon urination. Contact with the urine of maintenance hosts or contaminated soil and water is an opportunity for percutaneous infection of animals and humans, resulting in a range of symptoms, such as fever, headache, jaundice, and multi-organ failure (10, 12). Even limited to human cases, leptospirosis causes over 60,000 deaths annually worldwide, primarily in tropical regions (13), thus posing a public health problem requiring early address.

**Fig. 1.**
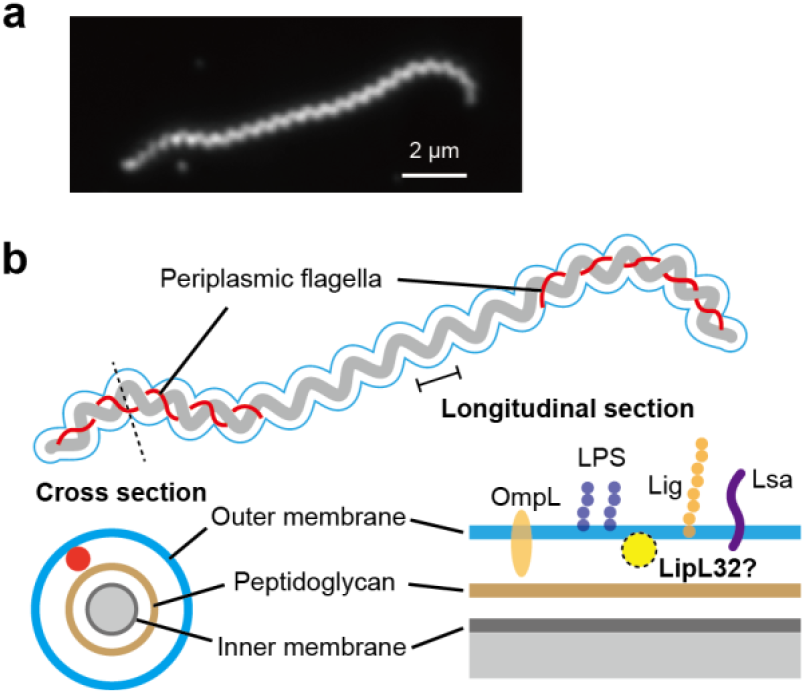
Morphology and structure of *Leptospira interrogans*. (a) A dark-field micrograph of *L. interrogans*. Periplasmic flagella curve the cell ends, and the entire cell body exhibits a short-pitch helix. (b) Schematic diagrams of the whole cell and membrane structures. Short periplasmic flagella (red) extend from both cell ends. The cross-section and longitudinal-section views illustrate the membrane structure typical of Gram-negative bacteria. Representative outer-membrane components are indicated in the longitudinal-section view. LipL32 is known not to be exposed to the outer-membrane surface, but its subcellular location is the subject of this study.

Vaccination is a key strategy for preventing the dissemination of leptospirosis; however, currently available vaccines are serovar-specific inactivated vaccines targeting LPS. Because *Leptospira* spp. are classified into more than 300 serovars based on LPS structure, the identification of broadly protective vaccine targets remains an important challenge. In addition to LPS, the OM of *Leptospira* spp. contains diverse protein components (Fig. 1b), some of which are conserved among pathogenic species and have therefore been investigated as candidates for the development of more broadly protective vaccines. For example, immunization by combining two OM proteins, LipL41 and OmpL1, provides a protective effect (14). Immunogenic properties of LigA and LigB, containing immunoglobu-lin-like repeats, are also confirmed in mice (15). LipL32, assayed in this study, is commonly carried by pathogenic species and accounts for ∼75% of the total OM protein (16). LipL32 is not exposed to the membrane surface (17), and immunization by the protein does not induce a protective immune response (18). Although the lipoprotein unique to pathogens has been expected to be a crucial virulence factor, the mutant lacking LipL32 retains pathogenicity, causing an acute manifestation comparable to that of the parental strain (19). Thus, despite being a pathogen-specific protein, LipL32 is not believed to be an essential virulence factor, and its role remains unclear.

In this study, we sought to elucidate the significance of LipL32 by determining its subcellular location *in situ* using cryo-electron microscopy and characterizing the effect of LipL32 deficiency on the bacterial physical properties using biophysical experiments. Our experiments revealed a novel mechanism by which an abundant but non-essential lipoprotein contributes to the cell envelope integrity and cell-body stiffness of *L. interrogans*. We also bridge the structural and mechanical functions of LipL32 to spirochete infection.

## Results

### LipL32 forms a thin layer and defines the periplasmic width

An immunofluorescent assay with an anti-LipL32 antibody showed that LipL32 was detected in *L. interrogans* cells whose OM was permeabilized with methanol (17). This demonstrated that LipL32 is not exposed on the OM surface, but the precise subcellular location of LipL32 in the leptospiral cell has been elusive. To identify the sub-cellular localization of LipL32 in the wild-type (WT) *L. interrogans*, we used a *lipL32*-disrupted mutant generated by random transposon insertion (*lipL32::*Tn, hereafter ΔLipL32) as a control (Fig. S1) and compared their membrane structures by cryo-electron microscopy (cryo-EM). Disruption of *lipL32* did not affect the growth rate and cell morphology observed by dark-field microscopy (Fig. S2), but cryo-EM showed irregular deformation in the OM (Fig. 2a). We found an apparent layered density in the inner part of OM facing PG in the WT cells, which vanished in the ΔLipL32 strain and was confirmed in its *lipL32*-complemented strain and the WT carrying an empty vector (Figs. 2b). This observation suggests that LipL32 is localized on the periplasmic side of the OM, possibly via insertion of its lipid moiety into the membrane, forming a thin layer immediately beneath the OM.

**Fig. 2.**
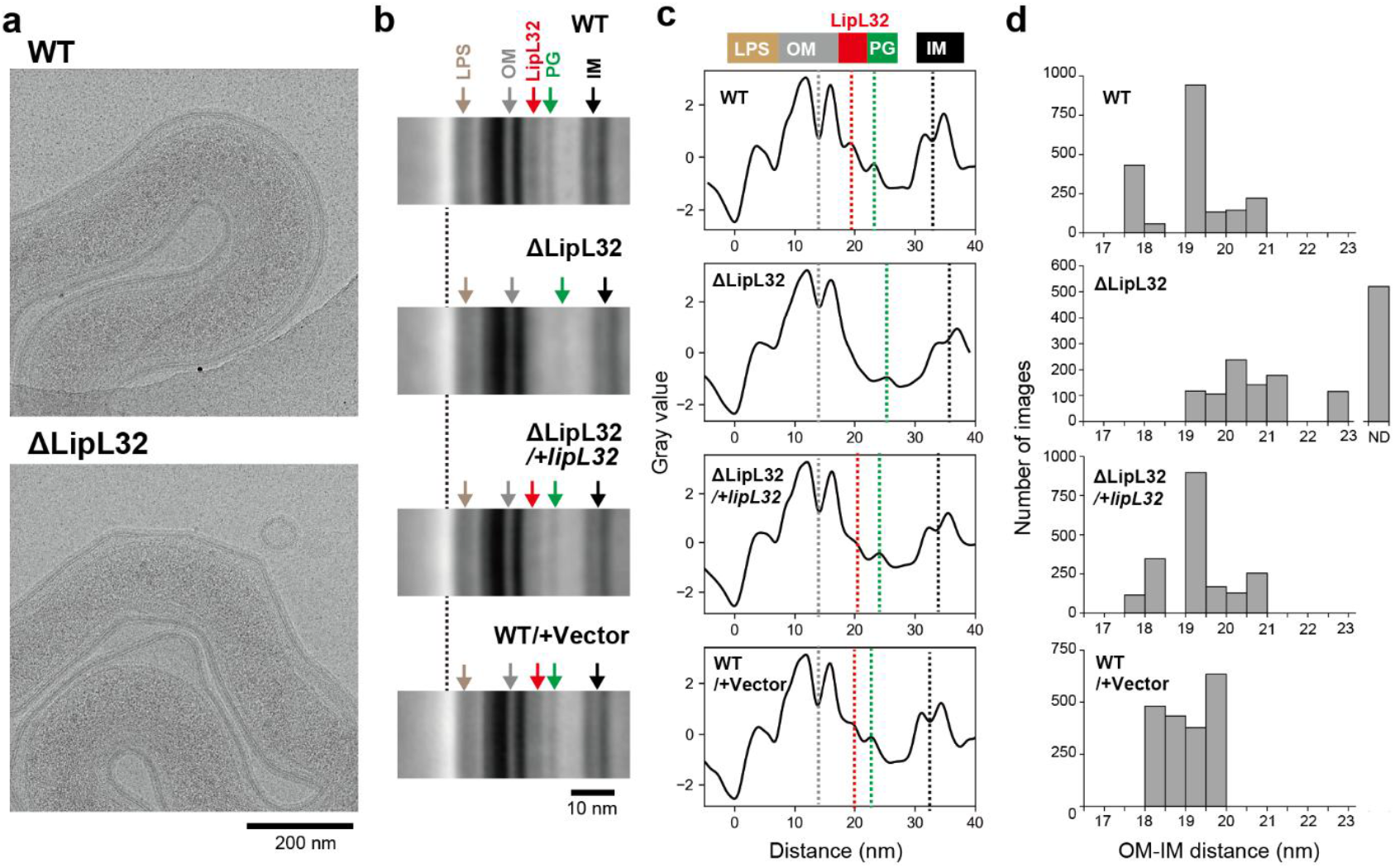
Effect of Δ*lipL32* on the membrane structure. (a) Cryo-electron micrographs of the WT and ΔLipL32 cells. (b) Enlarged average images of the cellular envelope comparing WT, ΔLipL32, the complemented strain (ΔLipL32/+*lipL32*), and the vector control (WT/+Vector). The images are aligned by matching the outer position of the LPS layer (vertical dashed lines). The membrane layers are indicated: outer membrane (OM), peptidoglycan layer (PG), and inner membrane (IM). Note the appearance of a distinct inner OM layer (red arrows) present only in the WT, the complemented strain, and the vector control. The results of 2D clustering analysis are shown in Fig. S3. (c) Averaged density profiles across the cell envelope. (d) Distribution of the OM-IM distance (periplasmic width). ND (No Data) category accounts for images where the IM was undetectable within the ROI (i.e., distance > 25 nm) or where IM density was averaged out during image averaging due to positional instability of IM.

We also found that the distance between the OM and IM of the ΔLipL32 mutant was wider than that of strains expressing LipL32 (Figs. 2c and d). Notably, the IM could not be detected in about 40% of the analyzed images of the ΔLipL32 mutant (represented as ND in Fig. 2d). In addition, both the position and density of the PG layer were also altered in the ΔLipL32 mutant (Figs. 2c and S4). These IM-undetectable images suggest that the IM is not fixed at a uniform position but varies between cells, leading to a reduction in signal intensity below the detection threshold due to image averaging, or that the IM is located beyond the measurable range (i.e., OM-IM distance > 25 nm).

### Cell stiffness

The structural evidence yielded by the cryo-EM implied the effect of *lipL32-*disruption on the mechanical properties of the cell membrane due to the loss of the LipL32-layer and the disruption of membrane integrity (Fig. 2). Therefore, we compared the rigidity of the WT and ΔLipL32 strains by measuring individual cells using optical tweezers. Among the cells adhering to the glass surface, cells whose bead-adhered cell ends were apart from the glass surface were selected, and the beads were trapped with an infrared laser (Fig. 3a). The cell was bent by moving the microscope stage to change the trapped bead position, and then the recovery of the bead position (cell morphology) to the original position after releasing from the laser trap was traced using machine-learning combined image analysis (Figs. 3b and S5, and Supplementary Information). The cell rigidity was estimated as a spring constant in this experiment, computed from the recovery rate of the bead position and drag coefficient acting on the bead (Fig. 3c). The stiffness measurement revealed that ΔLipL32 reduced the cell stiffness by half of the WT: 0.19 ± 0.02 pN/µm for WT and 0.11 ± 0.01 pN/µm for ΔLipL32. The reduced rigidity of ΔLipL32 was recovered by the complementation of the *lipL32* gene up to the WT level (0.18 ± 0.02 pN/µm). Thus, an anomaly in the cell-envelope integrity significantly reduces the cell strength.

**Fig. 3.**
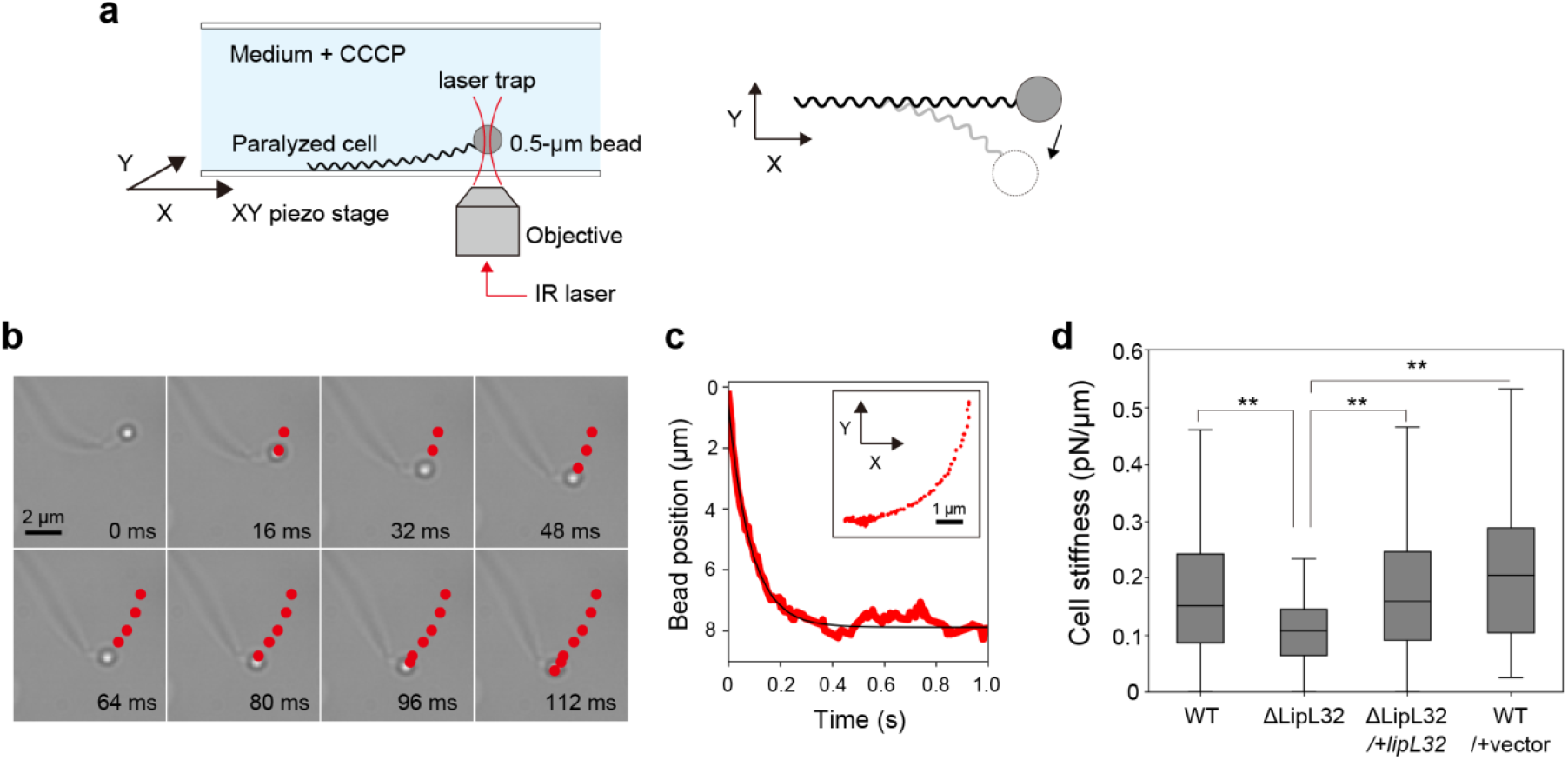
Effect of ΔLipL32 on the cell stiffness. (a) Schematic of the optical tweezers setup used for measuring cell stiffness (left panel). The right schematic illustrates cell bending by moving the piezo stage in the X-Y plane. (b) Example of bead position recovery after release from the laser trap. At 0 ms, the bead attached to the cell end is trapped by a laser. Red dots indicate the bead positions. The example shows two-dimensional movement, but 3D trajectory analysis was performed for all measurements (details in the Supplementary Information and Fig. S5). (c) Time-dependence of bead displacement after release (red line). The black line is the result of exponential fitting by *A*exp[(− *k*/*γ*)*t*] + *C*, where *k* is the spring constant (measured as a cell stiffness in this study), *γ* is the drag coefficient, and *A* and *C* are arbitrary constants included in the fitting function (see Materials and Methods for details). The trajectory of the bead positions is shown in the inset. (d) Cell stiffness values for WT (n = 60 cells), ΔLipL32 (n = 49 cells), ΔLipL32/+*lipL32* (n = 56 cells), and a vector control (n = 34 cells). Statistical analysis was performed by the Mann-Whitney U test (\*\**P* < 0.01).

### Penetration into a heterogeneous gel

The major portal of entry for *L. interrogans* is thought to be injured skin that reaches the dermis (10, 12). Given this infection route and our optical tweezer data, we hypothesized that reduced stiffness caused by the loss of LipL32 would impair penetration into host tissue. To test this, we compared the penetration ability of WT and ΔLipL32 mutant strains into agar gels with heterogeneous mesh structures resembling dermis (Fig. 4a). The substrate modulus of skin dermis is 20 – 50 kPa (20), which is a very similar range of 1 – 2% agar (21). The pore size of 1% agar is 100 – 300 nm (22), comparable to the *L. interrogans* cell size (∼150 nm in width), and it decreases with the agar concentration (23). Therefore, this *in vitro* assay allows us to qualitatively evaluate the mechanical performance of the *L. interrogans* penetration.

**Fig. 4.**
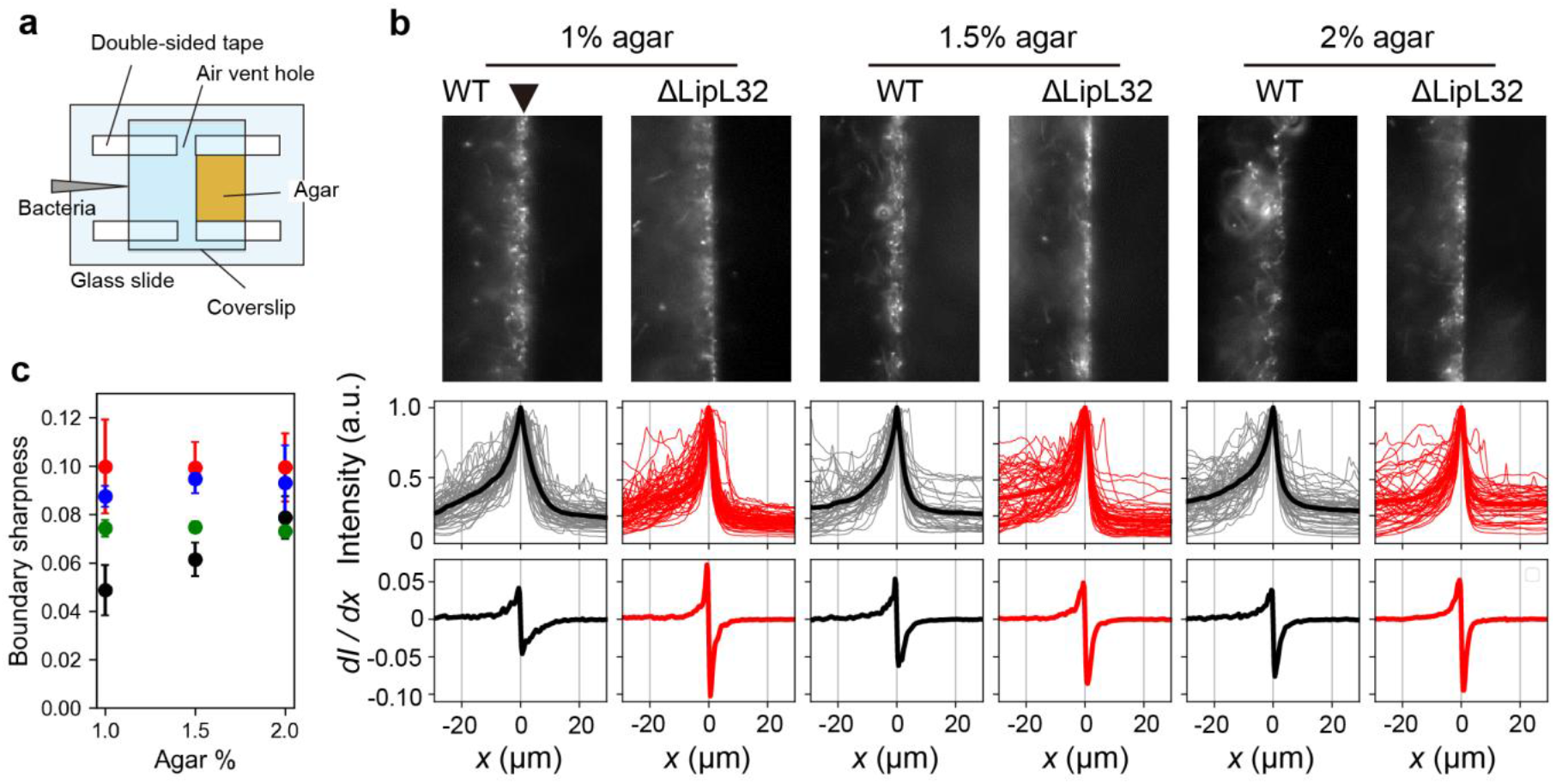
Bacterial penetration into a heterogeneous gel. (a) A flow chamber containing a liquid-agar boundary. (b, top) Micrographs of liquid-gel interfaces exposed with the WT and ΔLipL32 cells, examined in the presence of 1%, 1.5%, and 2% agar. The black triangle shown in the micrograph of WT in 1% agar indicates the boundary of the liquid containing bacteria and agar. The middle panels show brightness profiles around the liquid-gel boundaries (results of ΔLipL32/+*lipL32* and vector control are shown in Fig. S6). The spatial gradient of brightness (bottom), *dI*/*dx*, was determined by differentiating the brightness profiles. Larger values correspond to sharper boundaries. (c) Dependence of the boundary sharpness on the agar concentration: Black for WT, red for ΔLipL32, blue for ΔLipL32/+*lipL32*, and green for a vector control. The maximum values of *dI*/*dx* were defined as the boundary sharpness. Average values and standard deviations of three independent experiments are shown.

The penetration assay showed that WT cells disrupted the gel–liquid interface, creating rough boundaries due to bacterial entry. In contrast, ΔLipL32 mutants attached to the gel surface but failed to penetrate deeply, leaving a sharp liquid–gel interface intact (Fig. 4b). The sharpness of the liquid-agar boundary exposed to the WT cells was increased with the agar concentration, meaning impairment of bacterial penetration at high agar concentrations (Fig. 4c). These results suggest that a certain level of cell stiffness is critically required for efficient penetration into the dense tissue-mimicking matrix. (Results of the *lipL32*-complemented and vector-control strains are discussed below.)

## Discussion

This study sought to elucidate the enigmatic role of LipL32, the most abundant OM lipoprotein in the zoonotic spirochete *L. interrogans*. Despite its specific presence in pathogenic species, its precise function and necessity for virulence have been debated. Using cryo-EM and biophysical experiments, we have provided structural and functional evidence demonstrating that LipL32 forms a distinct layer in the periplasmic space that acts as a critical mechanical stabilizer for the cell envelope and confers strength on the cell body. The reduced cell stiffness by the loss of LipL32 affects penetration into tissue-mimicking matrices.

### “Lateral pressure” model based on high-density LipL32 packing

Our most striking finding is the cryo-EM visualization of a layered density formed by the highly abundant LipL32 immediately beneath the OM in the periplasmic space (Figs. 2b and c). Our results revealed an unexpected arrangement where this massive amount of protein is tightly packed to form a structural layer that is distinct from the conventional three-membrane structure of Gram-negative bacteria, OM, PG, and IM (Fig. 5a, upper panel). Although LipL32 molecules may lack the lateral-bonding characteristic of an actual biological membrane, their dense organization could function as a pseudo-structural scaffold stabilized by lateral packing pressure (black arrows in Fig. 5a, upper panel). While cryo-EM showed irregular deformation of the cell envelope upon *lipL32* disruption (Figs. 2a, lower panel), such structural defects were not observed by dark-field microscopy (Fig. S2). Most likely, the membranes of the ΔLipL32 mutant, which had been fragilized by the loss of LipL32 periplasmic layer, suffered more severe damage by external forces such as rapid water absorption during the cryo-EM sample preparation process than those of WT. This may explain the increased OM-IM distance observed by cryo-EM (Fig. 5a, lower panel), leading to the result that IM was not detected in nearly 40% of the ΔLipL32 strain data (Fig. 2d). Meanwhile, the fact that non-pathogenic *Leptospira* species synthesize PG normally without LipL32 (24) yields the hypothesis that the LipL32 layer is a pathogen-specific evolutionary enhancment to provide extra mechanical rigidity beyond that required for basic viability.

**Fig. 5.**
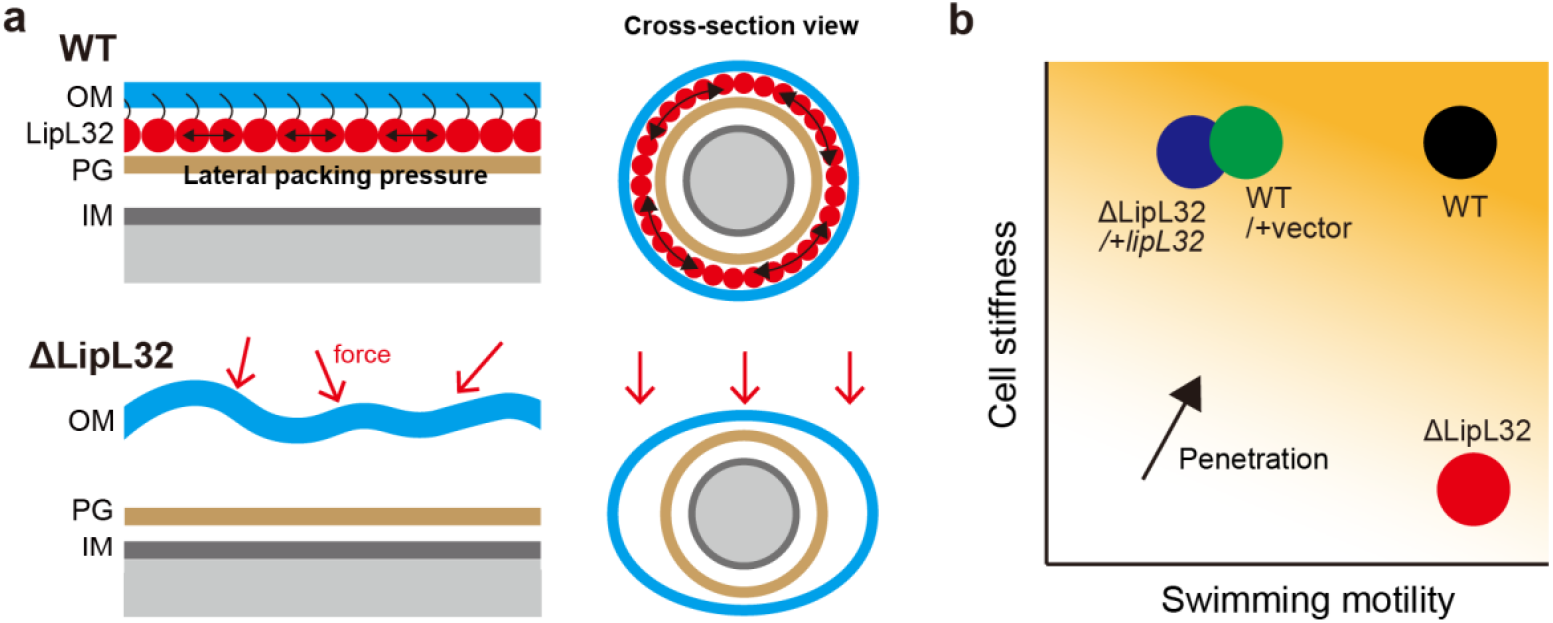
Proposed model for the structural role of LipL32 and association with the penetration ability. (a) LipL32 proteins (red spheres, upper panel) form a distinct, highly dense layer within the periplasmic space of *L. interrogans*. This dense packing is hypothesized to generate lateral packing pressure (black arrows), which acts as a structural scaffold to reinforce the outer membrane (OM). In the absence of the LipL32 layer (lower panel), the cell envelope becomes mechanically fragile and susceptible to external forces—such as the surface tension encountered during cryo-EM sample preparation—leading to the irregular membrane deformation and non-uniform periplasmic width observed in this study. The left schematics illustrate the cross-sectional view of the leptospiral cell body. (b) A reduction of the cell stiffness interferes with the bacterial penetration into gel-like structures, even while fully maintaining motility, vice versa. The thicker orange in the background represents higher penetration ability.

### “Tether and spring” model based on potential LipL32-PG interaction

While lateral packing pressure provides a feasible mechanism for OM reinforcement, the possibility of LipL32-PG interaction cannot be excluded, especially given that another OM lipoprotein, LipL21, is known to bind PG (25). Furthermore, elasticity measurement using an atomic-force microscope showed that the loss of a penicillin-binding protein (PBP) in *Staphylococcus aureus* reduces the cell rigidity (26). Although LipL32 lacks a typical PGB motif, assuming the direct interaction between LipL32 and PG allows for an alternative model. The structural demands of spirochetal membranes, which undergo constant cyclic deformation, evoke a functional similarity to eukaryotic axonemes (27). In the system, nexin linkers maintain the relative positioning of adjacent microtubule doublets and resist excessive shear deformation (28). By analogy, we hypothesize that the dense LipL32 layer functions as a structural tether that bridges the OM and PG, providing the necessary restoring force to stabilize the periplasmic architecture. High-density packing of LipL32 may realize multi-point physical interactions with the PG layer, acting as a mechanical spring that provides a restoring force against external stresses, such as the surface tension encountered during cryo-EM sample preparation. Whether and how LipL32 physically interacts with the PG layer remains an open question, and elucidating the underlying molecular mechanisms will be a compelling subject for future study.

### Significance of cell rigidity for penetration

As expected from the structural evidence, the loss of the LipL32 layer and the resulting periplasmic instability are translated directly into a significant loss of cell rigidity (Fig. 3). This demonstrates a direct link between the novel LipL32 layer and the overall mechanical robustness of the spirochete cell. The necessity of this rigidity was confirmed by the penetration assay (Fig. 4). The ΔlipL32 mutant, which is significantly softer, failed to penetrate the dense tissue-mimicking agar matrix, whereas WT cells effectively disrupted the gel interface. It has been suggested that the reduced cell rigidity of *S. aureus* due to PBP deficiency possibly affects the bacterial invasion into the osteocyte network (29). More broadly, cell mechanical properties are increasingly recognized as critical determinants of cellular function and invasiveness. For example, alterations in cell stiffness are known to impair capillary passage in malaria-infected red blood cells, highlighting the importance of appropriate mechanical flexibility for successful tissue traversal (30). Our results establish that, in addition to the known essentiality of motility, sufficient cell rigidity could be a critical and separable factor required for *L. interrogans* to penetrate dense, heterogeneous structures that mimic host dermal tissue.

### Interplay between cell rigidity and motility

While our primary data support the necessity of cell stiffness, the penetration assay revealed complexity: the complemented and vector control strains, despite exhibiting the WT-level stiffness (Fig. 3d), did not fully recover their penetration rates (Fig. 4c). As detailed in the Supplementary Information, we found that these mutants carrying plasmids slowed the swimming speed despite no impact on cell-body rotation (Fig. S7). The swimming-dependent locomotion of spirochetes, including *Leptospira*, is achieved by coordinated rotation of PFs residing at both ends of the cell body. The mechanism of PF coordination is elusive, but it is believed to involve the sensory and signal transduction, such as the chemotaxis system (5, 31). The genetic manipulation or antibiotics added to maintain the plasmids in the mutants might affect gene regulation and protein dynamics, associated with the coordinated PF rotation, independently of the structural role of LipL32. This observation is significant, as it suggests that both sufficient cell rigidity (mediated by LipL32) and efficient motility (mediated by PF coordination) are independent yet crucial determinants for optimal invasion to gel structure (Fig. 5b). This interpretation could reconcile our findings with previous animal studies that showed the ΔLipL32 mutant retains pathogenicity when inoculated into the eye and intra-peritoneally (19). We hypothesize that the structural role of LipL32, providing mechanical strength, is specifically essential for navigating the physical challenges posed by the dense extracellular matrix of the damaged skin dermis, which is the major natural route of infection. In contrast, infection via less physically restrictive routes (e.g., mucosal route such as the eye) may bypass the need for this enhanced rigidity.

In conclusion, our study defines LipL32 as a critical periplasmic structural element that provides the mechanical stability necessary for *L. interrogans* to breach the physical barriers of the host. This work identifies a novel mechanism by which an abundant but non-essential lipoprotein contributes to the cell mechanics of a pathogenic spirochete, establishing a new paradigm where cell rigidity is an infection route-dependent virulence factor. The elucidation of this specialized structural function is vital for understanding the pathogenesis of leptospirosis.

## Materials and Methods

### *Leptospira* strains

*L. interrogans* serovar Manilae strain UP-MMC-NIID (wild-type, WT) (15, 32, 33) the transposon insertion mutant *lipL32::*Tn (ΔLipL32), its complemented strain, and the WT strain carrying pNKLiG1 (vector control) (34) were used in this study. The ΔLipL32 mutant was generated by random *Himar1* mutagenesis of the WT strain via conjugation with *E. coli* β2163 harboring pCjTKS2, as described previously (35–37). The transposon insertion site was identified by semi-random PCR (37), revealing insertion between bp 85 and 86 of the 819-bp *lipL32* gene. The *lipL32*-complemented ΔLipL32 strain was constructed as follows. The *lipL32* gene, including its putative promoter region, was amplified from genomic DNA of the WT strain using PrimeSTAR GXL DNA polymerase (Takara Bio, Japan) with primers 5′-AAGCTTATCGATACCGTCGAGAACAAGAAAGAGTCAGAGAA-3′ and 5′-GCGAGGCTGGCCGGCGTCGATTACTTAGTCGCGTCAGAAGC-3′. The PCR product was assembled into *Sal*I-digested pNKLiG1 using the NEBuilder HiFi DNA Assembly kit (New England Biolabs, USA) and propagated in *E. coli* π1 (36). The resulting plasmid was transformed into *E. coli* β2163 and subsequently introduced into the *lipL32::*Tn strain via conjugation. *Leptospira* strains were cultured in liquid Ellinghausen–McCullough–Johnson–Harris (EMJH) medium (38) at 30ºC. The complemented and vector control strains were grown in EMJH medium supplemented with 25 µg/mL spectinomycin.

### Cryo-electron microscopy

Five milliliters of bacterial cultures were centrifuged at 1,000 × *g* for 5 min to concentrate the cells to a final volume of 50-100 µL. Aliquots of 2.5 µL were applied to glow-discharged holey carbon grids (Quantifoil R1.2/1.3, Au 300 mesh). The grids were blotted with filter paper for 2 s and immediately plunge-frozen in liquid ethane using a Vitrobot Mark IV (Thermo Fisher Scientific) operated at 4 °C and 100% humidity. Frozen grids were imaged using a Talos Arctica microscope (Thermo Fisher Scientific) operated at 200 kV and equipped with a Falcon 3 direct electron detector (Thermo Fisher Scientific). Dose-fractionated images were collected at nominal magnification of 92,000×, corresponding to a pixel size of 1.6 Å, with a total dose of 50 e^-^/Å^2^ and defocus values ranging from -0.6 to −1.4 μm.

All data processing was performed using RELION 5 (39). Movie frames were motion-corrected in RELION, and CTF parameters were estimated using CTFFIND4 (40). Particle picking was performed manually by tracing the membrane, which was treated as helical segments for particle extraction using a box size of 640 pixels with an inter-box distance of 150 Å. Approximately 4,000 particles were extracted and subjected to iterative 2D classification, each into 100 classes. Particles corresponding to segments in which the membrane was straight or convex toward the extracellular side were selected, and 2D classification was repeated until approximately 2,500 particles remained.

### Expression and localization of LipL32

Whole-cell lysates from 1.5 × 10^8^ leptospiral cells were prepared in SDS-PAGE sample buffer and separated on 5–20% SDS-PAGE gels, followed by Coomassie Brilliant Blue (CBB) staining. To examine the cellular localization of LipL32 in *L. interrogans* strains, cells from 12 mL cultures were extracted with Triton X-114, as described by Zuerner et al. (41), except that a protease inhibitor cocktail (Roche) was added during the extraction procedure. The resulting fractions were analyzed by 5–20% SDS–PAGE, followed by CBB staining.

### Growth assay

*Leptospira* cells grown in EMJH medium were diluted 1:10 into a fresh EMJH medium. EMJH media containing *Leptospira* cells were incubated at 30ºC, and their optical densities at 420 nm were measured.

### Morphology analysis using a dark-field microscope

*Leptospira* cells were harvested by centrifugation at 1000 × *g* for 10 minutes at room temperature. The bacterial precipitate was suspended in 20 mM potassium phosphate buffer (pH 7.4, PB) containing 100 µM CCCP (PB/CCCP), by which *Leptospira* cells were paralyzed (42). The paralyzed cells were infused into a flow chamber and observed with a dark-field microscope (microscope body BX50, oil-immersion objective × 100 LMPlanFLN, ×5 relay lens, oil dark-field condenser, high-brightness light source U-LGPS, Olympus, Japan). The cell images were captured by a CMOS camera (ORCA-Spark, Hamamatsu Photonics, Japan), and the morphology of individual cells was analyzed using ImageJ.

### Stiffness measurement

Cell stiffness was measured by optical tweezers constructed in our previous study (43). *Leptospira* cells were paralyzed as performed for the morphology analysis (see above). The paralyzed cells were infused into a flow chamber and incubated for 1 hour at room temperature. After flushing out floating cells with PB/CCCP, 1-μm polystyrene beads (ThermoFisher Scientific, Waltham, MA) diluted to 1:200 in PB/CCCP were infused into the flow chamber. The beads spontaneously attached to the bacterial surface without linkers (11) were confirmed, and floating beads were removed with PB/CCCP.

The optical-tweezers system (laser diode L1064H1, LD current controller LDC210C, Thorlabs Inc., Newton, NJ) was assembled into a bright-field microscope (microscope body IX70, oil-immersion objective ×100 LMPlanFLN, high-brightness light source U-LGPS, Olympus, Japan) and recorded using a CMOS camera (DFK33UX249, The Imaging Source, Germany) at a frame rate of 250 Hz. The optical tweezers trapped a bead adhering to a cell anchored to the glass. The cell body was bent by shifting the bead position using a piezoelectric stage. The bent cell body was recovered to its original shape by turning off the laser while resisting the viscoelastic force acting on the bead and cell body. The restoring force and drag force are given by *kR* and Ṙ respectively, where *k* is the spring constant of the cell body, *R* is the bead position, and *γ* is the drag coefficient of the sum of cell body (*γ*_C_) and bead (*γ*_B_): *γ*_C_ = 4*πµL*/[In(*L*/2*r*_C_) + 0.84] (44) and *γ*_B_ = 6*πµr*_B_ (from the Stokes’ law), where *L* in the cell length (measured for individual cells), *µ* is the medium viscosity (0.9 mPa ×s), *r*_C_ is the cell radius (0.07 μm), and *r*_B_ is the bead radius (0.5 μm). Assuming the balance between the restoring and drag forces yields *R* = *A*exp[(− *k*/*γ*)*t*] + *C* (see the Fig. 3 legend). Despite the introduction of a great approximation into the model, in agreement with this, tracking of the bead position after release from the trap gives a relaxation curve (Fig. 3c). The three-dimensional bead positions were traced by a custom-made image analysis program developed using Python (Supplementary information).

### Agar-penetration assay

The bacterial density of the cultured *Leptospira* was adjusted to 1 × 10^8^ bacteria/mL (OD_420_ ∼0.1) with EMJH medium. Agar (Bacto™ Agar) dissolved in PB was infused into a flow chamber and left at room temperature until solidified. The bacterial solution was infused into the chamber from the opposite side of the agar phase and incubated for 30 minutes at room temperature. The agar-liquid interface was recorded at a frame rate of 30 Hz, and the brightness of the area was analyzed with the ImageJ software.

### Motility assay

The *Leptospira* cells grown in EMJH were diluted 1:100 into PB and infused into a flow chamber. The bacteria were observed using a dark-field microscope (microscope body BX53, dry dark-field condenser, objective ×20 UPlan Fl, halogen lamp, Olympus, Japan) and recorded with a CMOS camera (ORCA-Fusion, Hamamatsu Photonics, Japan) at a frame rate of 100 Hz. Swimming speeds were measured by tracking the cell centroid as described in (45). The cell-body rotation was quantified by measuring the gyration of the hook-shaped cell end as described in (46).

## Acknowledgments

This work was supported by the JSPS KAKENHI: 21H02727 and 24K02274 for SN, and 22K07062 for NK.

## Author contributions

KA, NK, HK, MH, TK, and SN planned the project and wrote the manuscript. KA, NK, HT, and MH carried out the experiments and data analysis. KA and SN set up the optical system. HT, MH, and TK set up the electron microscope. KA, HK, MH, and TK made programs for data analysis. All authors contributed to the article and approved the submitted version.

## Competing interests

The authors declare that they have no competing interests.

## Data availability

The data supporting the findings of this study are available from the corresponding au-thor upon request.

## Notes

### Competing Interest Statement

The authors have declared no competing interest.

