## Supplementary Information and Figures for "A novel periplasmic layer formed by an outer membrane lipo-protein governs the cell-envelope integrity and stiffness of *Leptospira interrogans*"

Contents:

Fig. S1. Expression and localization of LipL32.

Fig. S2. Effect of ΔLipL32 on the morphology and growth.

Fig. S3. Flowchart and results of 2D cluster analysis.

Fig. S4. Effect of ΔLipL32 on the peptidoglycan layer (PG).

Fig. S5. 3D tracking of a bead attached to the *Leptospira* cell end

Fig. S6. Brightness profiles around the liquid-agar interface.

Fig. S7. Effect of ΔLipL32 on the *Leptospira* motility.

### Expression and localization of LipL32 in *Leptospira interrogans* strains

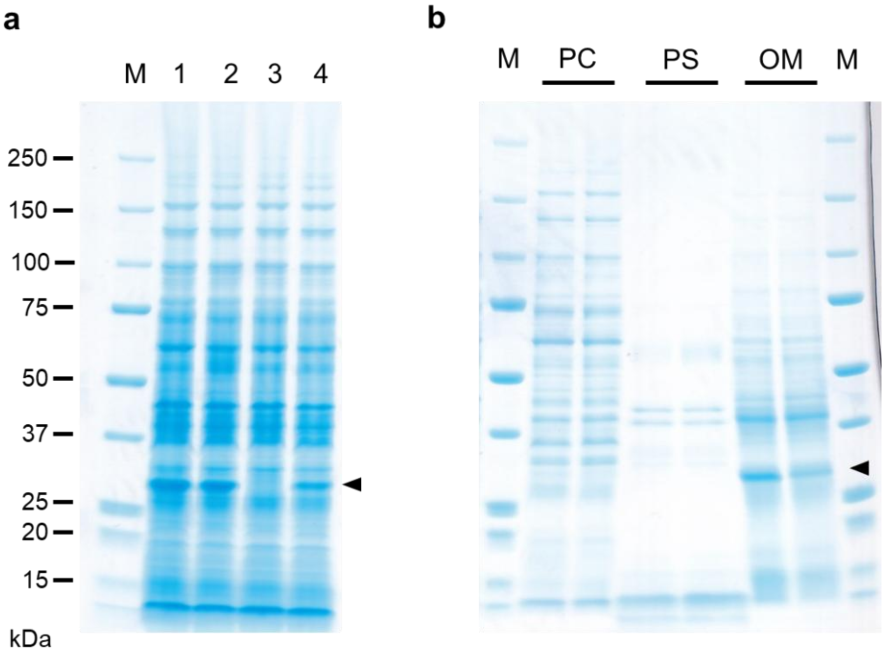


**Fig. S1. Expression and localization of LipL32.** (a) Expression of LipL32 in *Leptospira interrogans* strain UP-MMC-NIID: wild type (WT, lane 1), WT carrying an empty vector (lane 2), *lipL32*::Tn (ΔLipL32) mutant (lane 3), and *lipL32*-complemented ΔLipL32 mutant (lane 4). (b) Cellular localization of LipL32 in WT (left) and ΔLipL32 mutant (right). PC, protoplasmic cylinder; PS, periplasmic space; OM, outer membrane fractions. M, molecular weight marker. LipL32 is indicated by arrowheads. Experiments were performed independently twice, and each preparation was analyzed by 5–20% SDS-PAGE; representative gels are shown.

#### Effect of ΔLipL32 on the morphology and growth

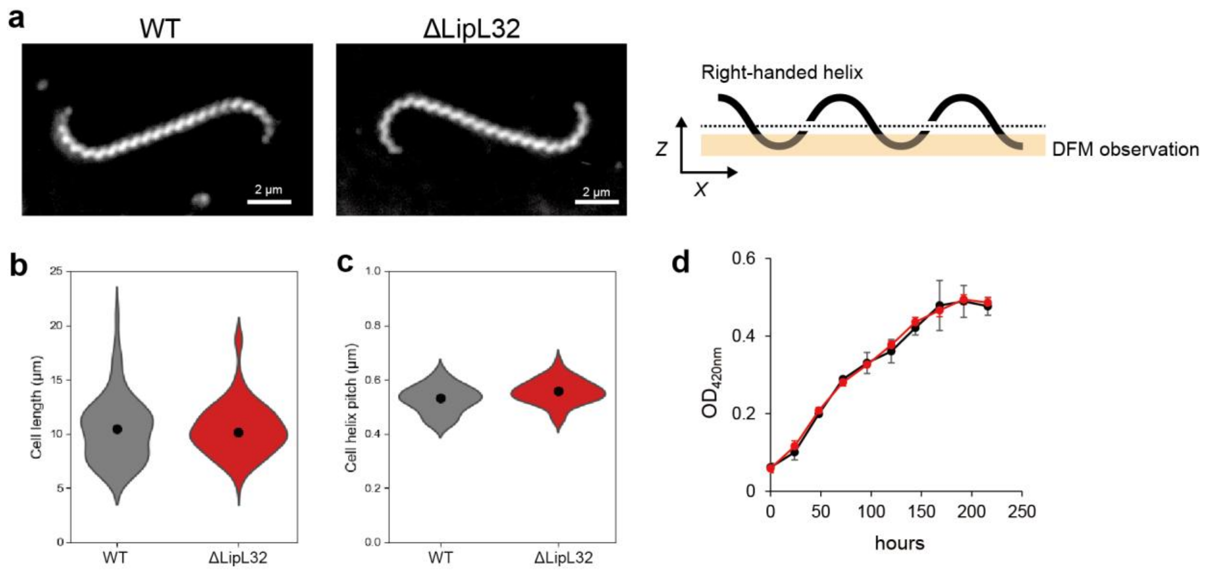


**Fig. S2. Effect of ΔLipL32 on the morphology and growth.** (a) Dark-field micrographs of WT and ΔLipL32 mutant of *L. interrogans* (left panels). Both strains were grown in EMJH liquid medium without antibiotics. The right panel explains how a right-handed helix of the leptospiral cell body is visualized using a dark-field microscope equipped with a high numerical aperture (NA) objective. A tilt observed along the cell body of WT and ΔLipL32 mutant indicates that both strains have a right-handed helical cell body. (b) Cell length obtained by measuring 66 WT and 101 ΔLipL32 cells. (c) Cell helix pitch obtained by measuring 63 WT and 70 ΔLipL32 cells for cell helix pitch. Black dots represent the median of each population in *b* and *c*. (d) Growth curves. The average values and standard deviations of three independent experiments are shown.

### 2D cluster analysis of cryo-EM data

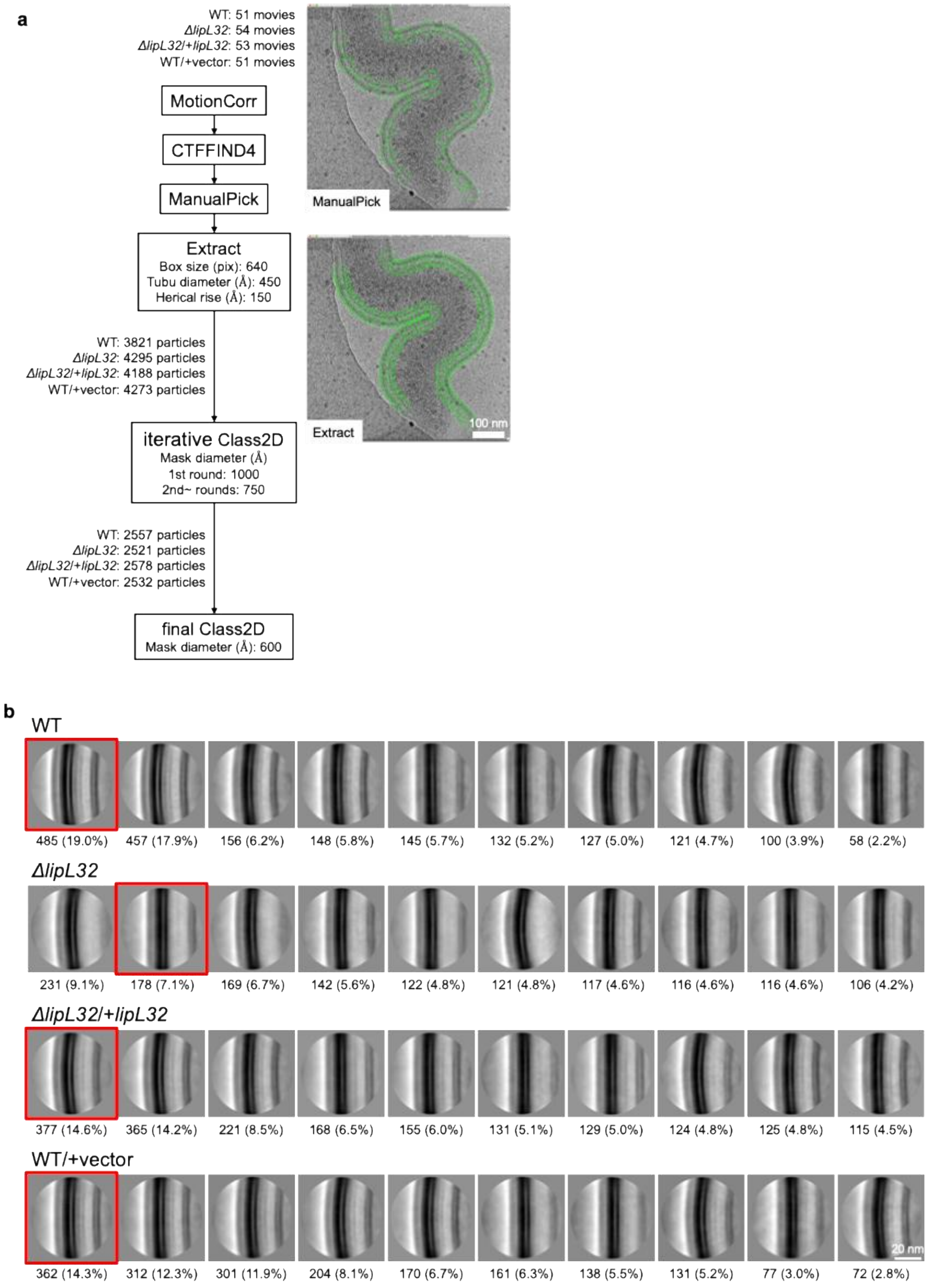

**Fig. S3. Flowchart and results of 2D cluster analysis.** (a) Flowchart of the cryo-EM analysis. (b) Top 10 2D class averages with the largest number of particles. Numbers below each class average indicate the number of particles contributing to that average. Percentages in parentheses represent the fraction of all particles used in the final 2D classification, with the total set defined as 100%. Images indicated by red boxes correspond to the original images shown in Fig. 2b.

#### Effect of ΔLipL32 on the peptidoglycan layer

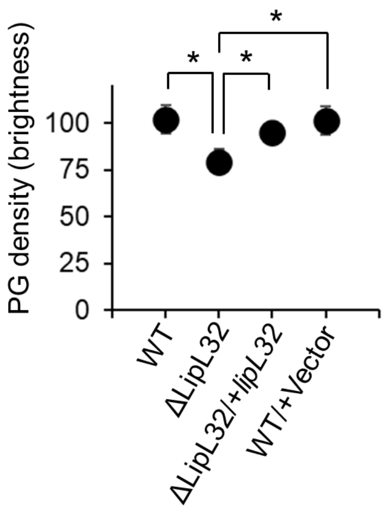


**Fig. S4. Effect of ΔLipL32 on the peptidoglycan layer (PG).** The brightness of the PG observed in sets of averaged Cryo-EM images (Fig. S3) was analyzed with ImageJ. The average values and standard deviations of the top 10 2D class averages are shown. The statistical significance between group averages was assessed by Weighted Least Squares (WLS) regression, considering the percentage of the total sample size used for each average as the statistical weight (*$P < 0.05$). Group comparisons were derived from the model coefficients. The analysis utilized the *statsmodels* library in Python.

#### 3D tracking of a bead attached to a bacterial cell

In the present cell-stiffness measurement, the cell body was bent by trapping and moving a bead attached to the cell end, and the bead position after releasing from the laser trap was tracked. Cells partially attached to the glass surface were targeted by optical tweezers, and bead movements usually show a three-dimensional (3D) trajectory. Regarding the 3D tracking, we solved several problems as described below.

**2D tracking.** The simplest two-dimensional (2D) particle tracking uses a threshold for discriminating the "target" particle from the background. This method is available when the brightness distribution of the target particle is almost constant, ideally. However, when we observe the recovery of a bead attached to the *Leptospira* cell end at the fixed Z-axis position of a bright-field microscope, the bead image changes with the Z-axis movement (Fig. S5a), making the threshold-dependent tracking difficult. To solve the problem, we first applied a machine-learning-based tracking, by which a dynamic foreground (a moving bead) is discriminated from a static background that is subtracted based on the Gaussian Mixture model (1). The machine-learning method allows us to perform 2D tracking of a moving bead. However, it cannot determine the bead position immediately before being released from the laser, because the background subtraction can recognize the static object (i.e., a trapped bead) as a background. Since the original position is essential for analyzing the recovery rate of the bead, we applied the conventional method for particle detection using an arbitrary threshold. That is, we performed 2D tracking by using the "threshold" method and the "machine learning" method for determining the original position and tracking a moving bead, respectively.

**Z-axis tracking.** To determine the Z-axis position at each time, we prepared a set of bead images with known Z-axis displacement from the focal plane as a reference (Fig. S5b). We determined the displacement in the Z direction by comparing the experimental data bead images, where the brightness distribution changes with displacement in the Z direction, with the reference based on the evaluation of the Structural Similarity Index Measure (SSIM) (Fig. S5c). Integration of 2D tracking and Z-axis tracking results yields a three-dimensional trajectory (Fig. S5d). Finally, the time-dependent displacement of the beads is determined from the three-dimensional trajectory, and the recovery rate is derived by fitting an exponential function (Fig. S5e).

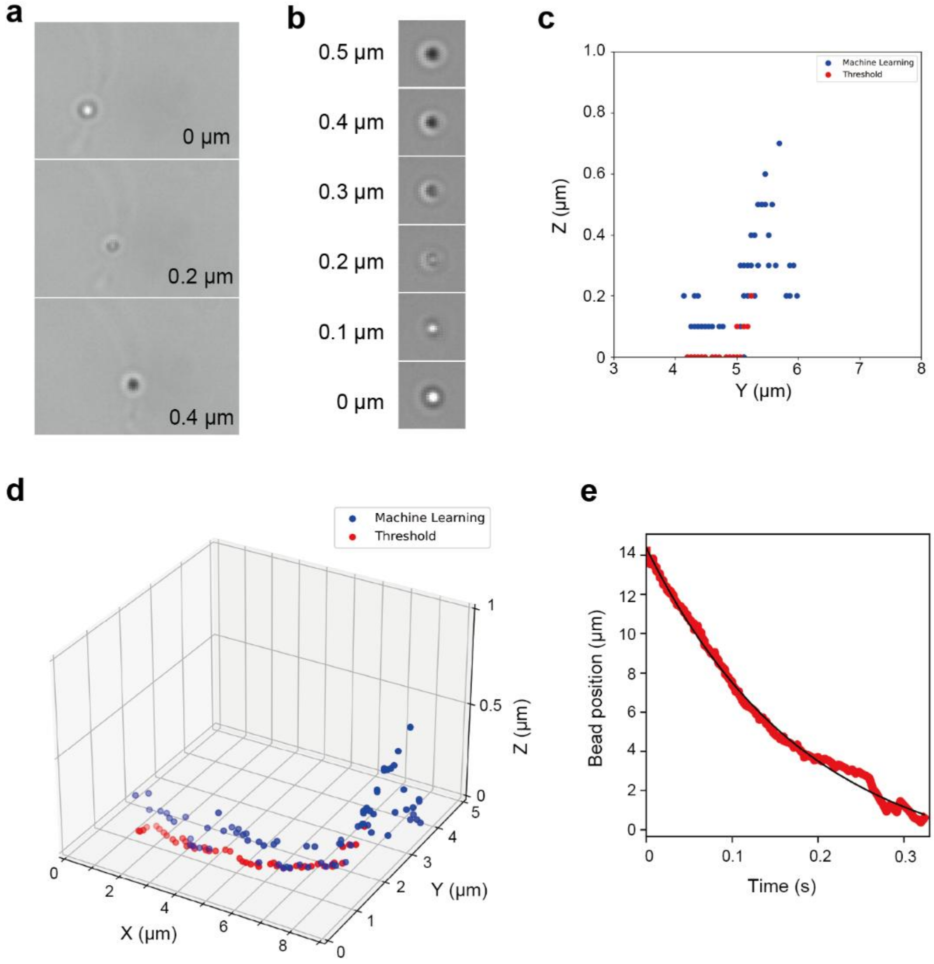


**Fig. S5. 3D tracking of a bead attached to the *Leptospira* cell end.** (a) Change in the brightness distribution of a bead moving with the recovery of the cell body after being released from the laser. (b) Bead images with known Z-axis displacement from the focal plane (0 μm). (c) Z-axis movement of a bead. The red and blue dots represent the results obtained by the "threshold" method and "machine learning" method, respectively. (d) A 3D trajectory. (e) A recovery curve obtained from *d*.

#### Bacterial penetration into agar

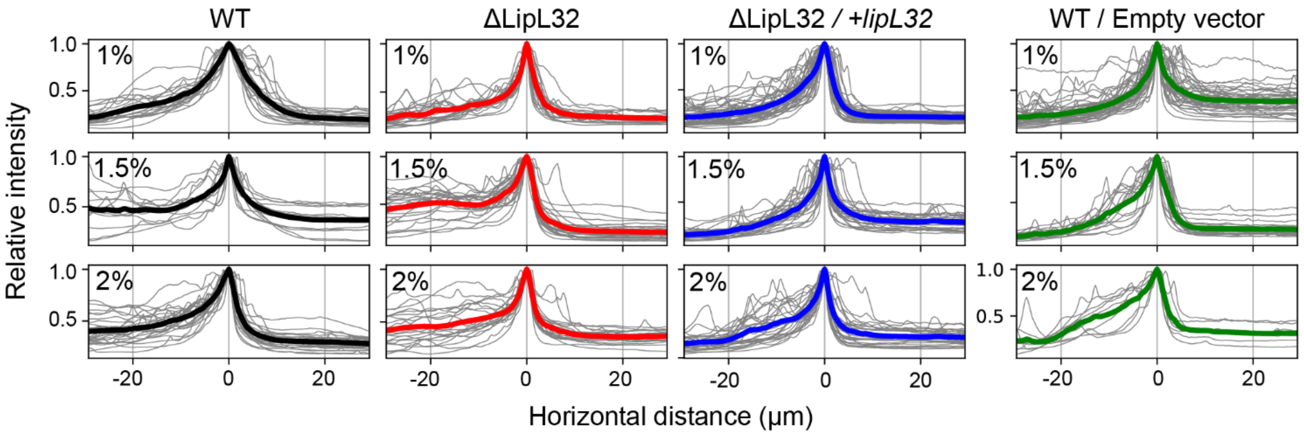


**Fig. S6. Brightness profiles around the liquid-agar interface.** Brightnesses of the liquid-agar interface (0 of the horizontal axis) exposed with the WT, ΔLipL32, ΔLipL32*/lipL32* (gene-complemented strain), WT/+Empty vector (vector control) cells were measured. The data of WT and ΔLipL32 are the same as those shown in the main text. The results obtained by three independent experiments are shown. See the main text for details of the experimental setup and analysis.

#### Effect of ΔLipL32 on the *Leptospira* motility

Regarding the insufficient recovery of gel-penetration observed in the *lipL32*-complemented strain and vector control, we assumed that genetic manipulations or antibiotic supplementation affected motility (see also *main text*). Motility is an essential virulence factor of *L. interrogans* (2, 3). The *Leptospira* motility form can be categorized into a "translational" mode, in which the cell is propelled forward by helical body rotation (Fig. S7a), and a "rotation" mode, in which the cell rotates without net displacement (Fig. S7b). Tracking of individual cells with distinguishing these two modes revealed that all strains exhibited similar levels of rotational activity (Fig. S7c), indicating that flagellar function was not impaired. However, the swimming speeds of the complemented and vector control strains were significantly slower than those of the WT and ΔLipL32 cells (Fig. S7d). Efficient swimming in spirochetes, including *Leptospira* spp., requires regulation of flagellar rotation at both cell ends, primarily mediated via chemotaxis signaling. Although the precise cause remains unclear, plasmid introduction or antibiotic selection may have affected intracellular signaling pathways to achieve smooth swimming.

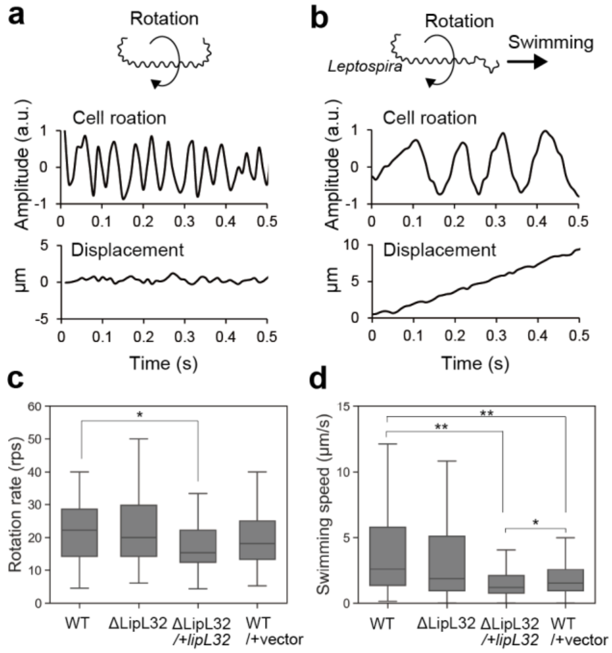


**Fig. S7. Effect of ΔLipL32 on the *Leptospira* motility.** (a) A schematic of the rotation mode (top), and example data of cell-body rotation (middle) and displacement obtained by measuring a typical rotation-mode cell (bottom). (b) A schematic of the translation mode (top), and example data of cell-body rotation (middle) and displacement obtained by measuring a typical translation-mode cell (bottom). (c) Rotation rates of the cell body. 57 cells for WT, 60 cells for ΔLipL32, 63 cells for ΔLipL32 /+*lipL32*, and 63 cells for a vector control were measured. Box plots show the median (horizontal line in the box), the 25th and 75th percentiles (box), and the minimum and maximum values within 1.5 times the interquartile range (whiskers). (d) Swimming speed. 145 cells for WT,138 cells for ΔLipL32,

148 cells for ΔLipL32 /+*lipL32*, and 169 cells for a vector control were measured. The box-and-whisker plots were made as in *c*. Statistical analysis was performed by the Mann-Whitney U test (*$P < 0.05$, **$P < 0.01$).
